# Proteomic assay for rapid characterization of *Staphylococcus aureus* antimicrobial resistance directly from blood cultures

**DOI:** 10.1101/2023.11.06.565783

**Authors:** Francis Deforet, Romain Carrière, Pierre L’Aour Dufour, Roxane Prat, Chloé Desbiolles, Noémie Cottin, Alicia Reuzeau, Olivier Dauwalder, Céline Dupieux-Chabert, Anne Tristan, Tiphaine Cecchini, Jérôme Lemoine, François Vandenesch

## Abstract

An efficient management of bloodstream infections requires a fast identification of the pathogen and a determination of its antimicrobial resistance profile. *Staphylococcus aureus* is among the most common pathogen causing bloodstream infection. A prompt characterization of methicillin-resistant *Staphylococcus aureus* (MRSA) and their aminoglycoside resistance profile is therefore crucial to quickly adapt the treatment and improve clinical outcomes. Among analytical technologies, targeted liquid chromatography-tandem mass spectrometry (LC-MS/MS) has emerged as a promising tool to detect resistance mechanisms in clinical samples. Herein we present a rapid proteomic workflow to detect and quantify the most clinically relevant antimicrobial resistance effectors in *S. aureus*: PBP2a, PBP2c, APH(3’)-III, ANT(4’)-I, and AAC(6’)-APH(2’’), directly from positive blood cultures and in less than 70 minutes. This approach provided 99% sensitivity for PBP2a (n=98/99 strains) detection. Sensitivity was 100% for PBP2c (n=5/5), APH(3’)-III (n=16/16) and ANT(4’)-I (n=20/20), and 94% for AAC(6’)-APH(2’’) (n=16/17). Across the entire collection, 100% specificity was reported for each of the 5 resistance proteins. Additionally, relative quantification of ANT(4’)-I expression allowed to discriminate kanamycin-susceptible and -resistant strains, in strains all harboring the *ant(4’)-Ia* gene. The LC-MS/MS method presented herein demonstrates its ability to provide a reliable and in-depth profiling of *S. aureus* resistance, directly from positive blood culture and in a short turnaround time, as required in clinical laboratories.

## Introduction

*Staphylococcus aureus* is one of the leading gram-positive bacteria involved in bloodstream infection (BSI). First line antibiotics used in *S. aureus* BSI are beta-lactam antibiotics targeting penicillin-binding proteins (PBPs), which are involved in cell wall synthesis. Methicillin-resistant *Staphylococcus aureus* (MRSA) strains are characterized by the expression of alternative PBP, PBP2a or PBP2c that have a reduced affinity to beta-lactams (1–4). These proteins are encoded respectively by the *mecA* and *mecC* genes carried by a mobile genetic element, the Staphylococcal cassette chromosome *mec* (SCC*mec*). There are 13 SCC*mec* types described to date, differing by the genetic organization of regulatory elements around the *mecA* gene and the type of recombinases (*ccr* genes) associated with the cassette (5). MRSA strains are of serious concern as they have been shown to be associated with higher mortality, increased hospital length of stay and increased medical costs (6–8).

Aminoglycosides, especially gentamycin and tobramycin are a second class of antibiotics relevant for the treatment of *S.aureus* BSI, especially in case of severe sepsis and/or prosthetic valve infective endocarditis (9, 10). Resistance toward these molecules are caused by the acquisition of aminoglycoside modifying enzymes (AMEs), which can inactivate antibiotics through the addition of a chemical group to the aminoglycoside backbone. In *S. aureus*, the most important AMEs are: (i) the 3’-O-phosphotransferase III (APH(3’)-III), encoded by the *aph(3’)-IIIa* gene, inactivates kanamycin and amikacin; (ii) the 4’-Oadenyltransferase I (ANT(4’)-I), encoded by the *ant(4’)-Ia* gene, inactivates kanamycin, neomycin, tobramycin and amikacin; and (iii) the 6’-N-acetyltransferase-2"O-phosphotransferase (AAC(6’)-APH(2")), encoded by the *aac(6’)-Ie-aph(2")-Ia* gene, confers resistance to gentamicin, tobramycin, kanamycin, amikacin and neomycin (11–15).

The positive impact of rapid antibiotic susceptibility testing (AST) and prompt administration of appropriate antibiotic therapy on the outcome of patients who experienced BSI or septic shock has been widely documented (16–18). Providing a solution that can offer reliable profiling of beta-lactam and aminoglycoside resistance in *S. aureus* from positive blood culture, with a short turnaround time and at reasonable cost is therefore of prime interest to improve *S. aureus* BSI management. Various molecular tests provide the presence/absence of the *mecA* and *mecC* genes in a short time (Xpert MRSA/SA BC by Cepheid, Biofire BCID2 by bioMerieux, ePlex BCID-GP Panel by GenMark, BD Max by BD, among others…) (19). Some others like the Unyvero system by Curetis also enable the detection of the aminoglycoside resistance genes *aac(6 ′)/aph(2 ′ ′)* and *aacA4* in about 5 hours (20). However, the relatively high cost per test of those molecular panels discourages clinician to implement it on a systematic basis. Also, despite being the gold standard method for identification of MRSA through detection of *mecA/C* genes, they do not provide expression levels of other resistance proteins. This information can be key for some resistance mechanisms, especially aminoglycosides, where the clinician is guided by the minimal inhibitory concentration (MIC), which depends on the expression of resistance genes. Today, there is no phenotypic method available to determine the antibiotic resistance from a positive blood culture in less than two hours. Among promising technological alternatives to true phenotypic methods, liquid chromatography coupled to targeted mass spectrometry (LC-MS/MS) approaches have been developed to reliably detect various classes of resistance proteins; these include carbapenemases, extended spectrum beta lactamases, porins, efflux systems, PBP2a, PBP2c, or proteins involved in aminoglycosides resistance (21–30). As the detection is performed at the protein level, LC-MS/MS can quantify the expression of the resistance effector, which is in some cases directly correlated with resistance levels (30, 31).

In the present study, a rapid targeted LC-MS/MS assay has been developed for the detection of *S. aureus* most relevant resistance mechanisms to beta-lactams (PBP2a and PBP2c) and aminoglycosides (APH(3’)-III, ANT4(‘)-I and AAC(6’)-APH(2’’)) in less than 70 minutes, directly from positive blood cultures.

## Materials and Methods

### Bacterial strains

A total of 104 MRSA and 20 methicillin susceptible *S. aureus* (MSSA) strains were selected in the collection of the French National Reference Center for Staphylococci (French NRCS) to account for the worldwide diversity of SCC*mec* types and strain genetic backgrounds (Supplementary Table S1).

### Peptides selection

In a former study by Charretier *et al.*, 8 PBP2a and 4 PBP2c peptides were identified as reliable surrogates for methicillin resistance detection in *S. aureus* cultured on agar plates (29). Because matrix effect and interferences can significantly vary between sample types (e.g. between bacterial colonies in suspension *versus* bacteria isolated from blood cultures), PBP2a peptides were tested for MRSA characterization in spiked blood cultures as described hereunder. Transitions choice was optimized, and the 4 best PPB2a peptides were included in the assay along with the 4 PBP2c peptides (Table S2).

For aminoglycoside resistance proteins, namely APH(3’)-III, ANT(4’)-I and AAC(6’)-APH(2’’) and, the protein sequences were digested *in silico* with trypsin using Skyline software (v20.2) to generate every potential surrogate peptides (32). Cysteine-containing peptides were excluded from the assay since the sample preparation protocol used in this study did not include steps disulfide bonds reduction and alkylation during the protein digestion.

All remaining peptides between 6 and 25 amino acids were kept to build a multiple-reaction monitoring (MRM) method containing 2+ and 3+ charge states and 3 y ions. Mass over charge values, declustering potential and collision energy were all predicted using Skyline software prediction equations for QTRAP6500 triple quadrupoles. This MRM method was then used to assess the detectability of each peptide by analyzing bacterial suspensions of strains carrying the corresponding resistance genes, according to Whole Genome Sequencing (WGS). Negative strains were also analyzed to confirm signal specificity. The peptides providing the most intense signal were then purchased (ThermoFischer Scientific, Rockford, IL, USA) as synthetic to confirm retention times and fragmentation patterns. In order to prevent any potential false positive and false negative results, variant coverage (i.e. the tryptic peptide is carried by all variants of the corresponding protein) was verified and sequence uniqueness was checked using Unipept 4.0 (https://unipept.ugent.be).

In addition, data-dependent analyses were performed externally on a Qexactive HF orbitrap (ThermoFisher) to identify *S. aureus*-specific ribosomal peptides to be used as controls for cell lysis and protein digestion rate, as well as markers for species confirmation. Their specificity was verified using BLAST searching and 6 ribosomal peptides were added to the MRM assay (Table S4). Signal specificity was confirmed with the corresponding synthetic peptides.

Also, ribosomal proteins being conserved and constitutively expressed in every *S. aureus* strains, their intensity can be used to normalize the levels of resistance peptides and obtain relative quantification values similarly to what is performed in RT-PCR experiments with housekeeping genes (30, 33).

To avoid quantification bias caused by potential methionine oxidation, only best flyers without methionine were used to perform relative quantification. Finally, two quality control peptides (AIVGASLDLIK and TGQSSLVPALTDFVR) were spiked in samples before LC-MS/MS analysis, to assess robustness and repeatability of the LC separation and MS detection across the whole duration of the study. Two peptides derived from auto-digestion of trypsin were also monitored to be used as controls of the protein digestion step.

### Blood cultures

*S. aureus* strains were grown on Columbia agar supplemented with 5% sheep blood (COS) (bioMérieux, Marcy l’Etoile, France) before blood culture. Aerobic and anaerobic blood culture bottles (BACT/ALERT FA/FN Plus®, bioMérieux) were spiked with 5 mL sterile human blood (obtained from the Etablissement Français du Sang) and approximately 1-10 bacterial cells in 1 mL of 0.45% NaCl. Bottles were incubated in a BACT/ALERT® VIRTUO® (bioMérieux) until flagged positive by the instrument. When not processed immediately, bottles were stored at 4°C for a maximum of 4 hours until sample preparation.

### Bacterial isolation and LC-MS/MS sample preparation

After positivity, 1 mL of blood culture was drawn from the bottle and transferred in a 1.5 mL tube. A total of 200 µL of a 6% sodium dodecyl sulfate solution was added, and the tube was mixed for 10 seconds, before a 2 minutes centrifugation at 16100g. The supernatant was discarded and the pellet resuspended in 1 mL LC-MS/MS-grade water (ThermoFisher Scientific). The tube was centrifuged 1 minute at 16100g, the supernatant discarded, and the pellet resuspended in 1 mL LC-MS/MS grade water before being transferred into a new 1.5 mL tube. Then, the tube was centrifuged again (1 minute, 16100g) and the supernatant discarded. Next, 200 µL of LC-MS/MS-grade water was added to the pellet as well as approximately 70 mg of glass beads (acid washed, 150-212µm, SigmaAldrich, St Louis, MO, USA). Bacterial lysis and protein digestion were performed simultaneously by adding 50 µL of 1 mg/mL trypsin (Roche Diagnostics, Mannheim, Germany) prepared in a 150 mM NH_4_HCO_3_ solution. The mixture was then placed in a thermostated (50°C) ultrasonic bath (Bioruptor Plus, Diagenode, Liège, Belgium); ultrasounds were applied at low power for 10 cycles (30 seconds on – 30 seconds off). Trypsin digestion was then stopped by adding 5 µL of formic acid. Digests were finally centrifuged for 5 minutes at 9600g and 100 µL of supernatant was transferred into the final screw cap vial. The total sample preparation time was 20 – 25 minutes from the positive blood culture to the peptide mixture.

### LC-MS/MS analysis

Samples were analyzed using an Agilent 1290 Infinity liquid chromatography (Agilent technologies) coupled to a QTRAP6500+ Triple-quadrupole mass spectrometer (Sciex, Toronto, Canada) equipped with an ESI Turbo V ion source. The instrument was operated in MRM mode. Mobile phases were H_2_O + 0.1% formic acid (Buffer A) and acetonitrile + 0.1% formic acid (Buffer B). The liquid chromatography separation was carried out on a Xbridge Peptide BEH C18 (1 mm × 100 mm, particle size 3.5 µm) column (Waters, Milford, MA, USA) heated at 60°C. Evaluation of PBP2a detection sensitivity was performed with a 15.5-minute gradient from 2% to 10% buffer B from 0 to 0.5 minutes and 10% to 40% buffer B from 0.1 to 15.5 minutes at a 75 µL/mL flow rate, 5 µl of sample were injected on the system. In order to fall under a 10-minute analysis turnaround time, development of the induction protocol and the final detection of resistance proteins in 124 *S. aureus* strains was performed using an optimized gradient; it consisted in 2% to 10% buffer B from 0 to 0.1 minutes and 10% to 35% buffer B from 0.1 to 5.95 minutes at a 100 µL/mL flow rate, for a total analysis time of 9.5 minutes. A total of 10 µL were injected on the system. For each peptide, m/z values, declustering potential and collision energy were obtained using Skyline software. Transitions lists used in this study are available along with raw data on PASSEL with the accession number PASS05841.

### Whole genome sequencing

Illumina libraries were prepared using the Nextera XT or DNA Prep kits (Illumina, San Diego, CA, USA) and sequenced on either a MiSeq or NextSeq 550 instrument (Illumina) using a 300 or 150 bp paired-end protocol, respectively. Assemblies were produced using SPAdes v3.14 and assembled genomes were used to perform MLST (https://pubmlst.org) typing, SCC*mec* typing (SCCmecFinder) and assessment of *in silico* resistance profiles (ResFinder database v.2021-03-0).

### Phenotypic susceptibility testing

AST was performed using broth microdilution (Sensititre, ThermoFisher). MICs were determined for the following antibiotics: cefoxitin, oxacillin + NaCl 2%, ceftaroline, ceftobiprole, kanamycin, tobramycin, gentamicin, erythromycin, clindamycin, quinupristin-dalfopristin, ofloxacin, tetracyclin, trimethoprim/sulfamethoxazole, rifampicin, fusidic acid, linezolid, tedizolid, vancomycin, teicoplanin, daptomycin, dalbavancin, and muropicin. The results were interpreted according to the 2020 guidelines of the French Committee for Antimicrobial Susceptibility Testing.

### Protocol optimization – PBP2a induction

Upon positivity, 3.8 mL of blood culture medium were drawn from the blood culture bottle and transferred to a 15 mL tube. For each condition, 200 µL of inducer was added. The different inducers were tested at following final concentrations: 0.4 µg/mL cefoxitin (Sigma-Aldrich), 5 mg/mL 6-animopenicillanic acid (6-APA) (Sigma-Aldrich), 5 mg/mL 7-aminocephalosporanic acid (7-ACA) (Sigma-Aldrich), 5 µg/mL clavulanic acid (Sigma-Aldrich), 1.5 mg/mL serine hydroxamate (Sigma-Aldrich), 5 mM H_2_O_2_ (Sigma-Aldrich), 0.1 mM methyl viologen (Sigma-Aldrich), 0.25 mM dipyridyl (Sigma-Aldrich), 0.03 µg/mL mupirocin (Sigma-Aldrich), 0.5 mg/mL vancomycin (Mylan), 5 mM H_2_O_2_ (Sigma-Aldrich), 5% NaCl (Sigma-Aldrich), or incubated at 30°C. Tubes were then incubated for 1 h, at 37°C under agitation (180 rpm). Cefoxitin and 6-APA induction were also tested for 10 minutes, 30 minutes, 1 hour, 2 hours, and 4 hours. Finally, 1 mL of induced blood culture medium was transferred to a 1.5 mL Eppendorf tube for bacterial isolation and sample preparation as described in “Bacterial isolation and LC-MS/MS sample preparation”.

### Peak detection and relative quantification

Raw chromatograms were analyzed using SciexOS 3.0 software for automatic peak detection and surface integration (autopeak algorithm) using low smoothing, a retention time half window of 30 seconds and the interference resolution parameter set at 50%. For each transition, peak integration was manually checked and curated when necessary to ensure a correct quantification. Label-free quantification strategies have now been widely applied, and demonstrated their robustness for quantification of proteins in biological samples (30, 31, 33). These approaches rely on the assumption that the signal of the most intense transitions of the best flying peptide correlates with protein abundance and can therefore be used as indicator of protein amounts (34). Herein, the area of the three transitions of quantifier peptides were summed to compensate for potential slight variations in fragmentation repeatability across runs. Also, to consider bacterial load variations across samples, signals of resistance quantifier peptides were expressed relative to signals of three housekeeping ribosomal peptides (SQSVLVFAK, VLELVGVGYR and ILEEANVSADTR) derived from ribosomal proteins that were used as an indicator of bacterial density due to their quantotypic properties (30, 33). Peptides used as quantifiers are reported in table S4. In other words, quantification values are obtained using Equation 1:

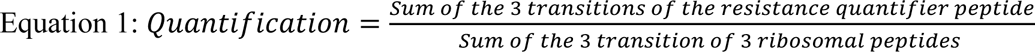

To note, PBP2a relative quantification results presented in Figure 1 were obtained by normalizing the signal of the PBP2a quantifier peptide ELSISEDYIK with the signal of the *S. aureus* ribosomal peptide SQSVLVFAK only, because a different MS method was used at that time.

**Figure 1.**
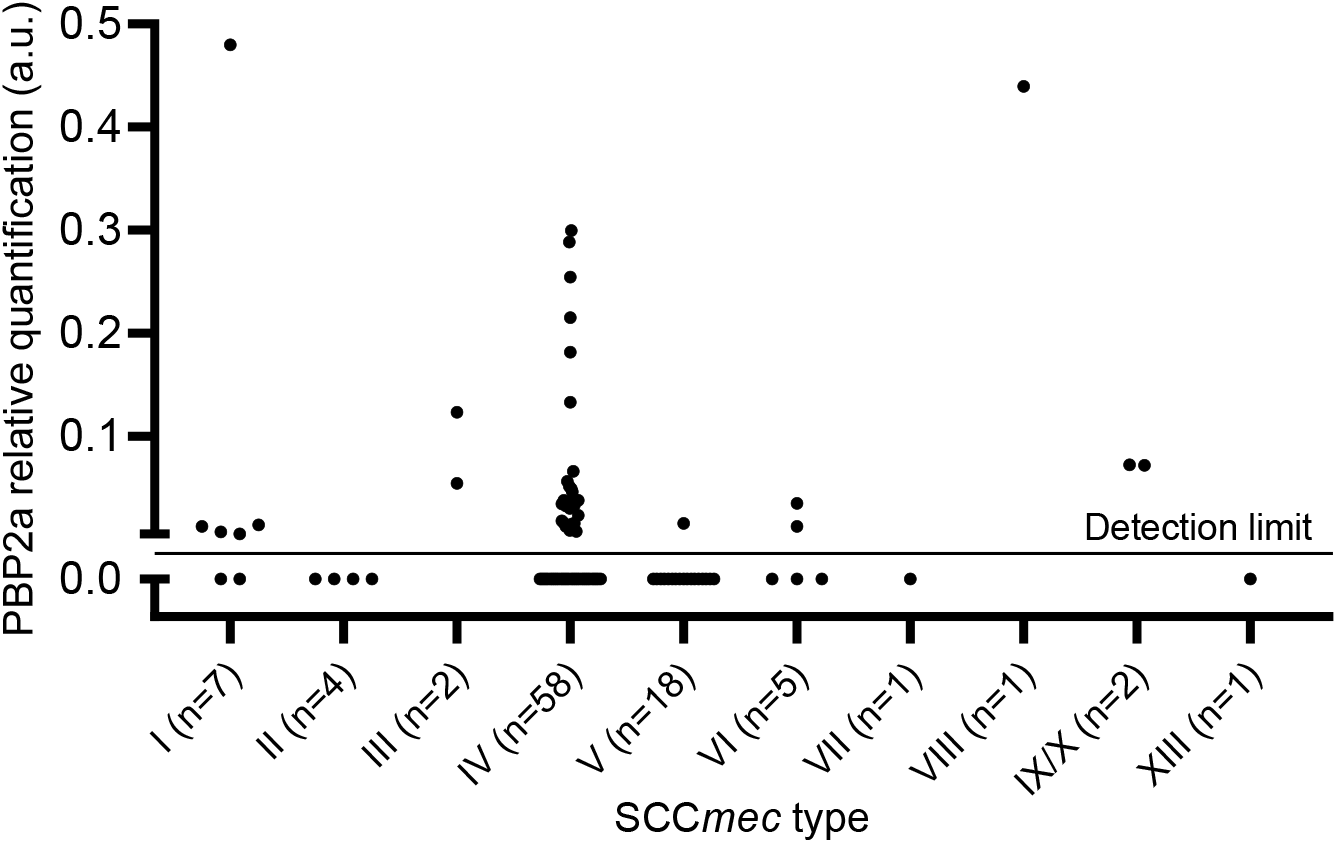
Relative quantification of PBP2a expression in 99 mecA MRSA strains with various SCCmec elements, representative of the worldwide diversity. Signal of peptide ELSISEDYIK was normalized on the signal of the ribosomal peptide SQSVLVFAK. When the three transitions of ELSISEDYIK were not correctly detected, quantification was set to zero.

### Statistical analysis

Statistical analysis presented in figure 5 was performed using Mann-Whitney U test. Differences between PBP2a expression levels upon induction with cefoxitin, cefoxitin + mupirocin and 6-APA were performed using a Student t-test. Differences were considered significant for p-values < 0.05.

## Results

### Sensitivity evaluation of MRSA detection

Sensitivity regarding PBP2c detection in *mecC* MRSA was not evaluated due to the low number of isolates at our disposal. To evaluate the sensitivity regarding PBP2a detection in MRSA blood cultures, LC-MS/MS analysis was carried out on 99 *mecA* positive MRSA and 20 MSSA strains of the French NRC.

Validation criteria for PBP2a detection was set to detection of at least 3 transitions of peptide ELSISEDYIK. The manual analysis of the chromatograms confirmed the detection of PBP2a in only 43 out of 99 *mecA* positive MRSA strains (43.4% sensitivity). No signal was observed in any MSSA samples. Relative quantification revealed a broad heterogeneity of PBP2a expression among PBP2a positive samples (figure 1).

PBP2a was correctly detected for only 28/58 (48.2%) type IV SCC*mec* strains. Among PBP2a-positive type IV SCC*mec* strains, an up to 40-fold difference in PBP2a relative expression level was observed. Besides, PBP2a detection was validated in only 1/18 (5.5%) strains with type V SCC*mec*, illustrating the impact of the *mecA* genetic environment on *mecA* expression and the underlying regulation dynamics.

### Protocol optimization – PBP2a induction

The 43.4% sensitivity performance is unacceptable for any fit-for-purpose MRSA diagnostic tool, therefore, a workaround strategy involving *mecA* expression induction was implemented. PBP2a expression in *S. aureus* is regulated and can be activated by diverse mechanisms such as beta-lactam exposure (35, 36), stringent stress (37, 38), oxidative stress (39, 40), osmotic stress (41), temperature (39) or glycopeptide exposure (42). Thus, 13 different treatments were tested for their ability to trigger PBP2a expression (figure 2), while minimizing the impact of this induction step on the global turnaround time. Among the MRSA strains in which PBP2a was not detected in the previous results, three strains (STAAUR35, STAAUR64 and STAAUR93) were used to set up the induction protocol. These 3 strains were selected as most of false negative MRSA harbored SCC*mec* types II, IV, or V.

**Figure 2.**
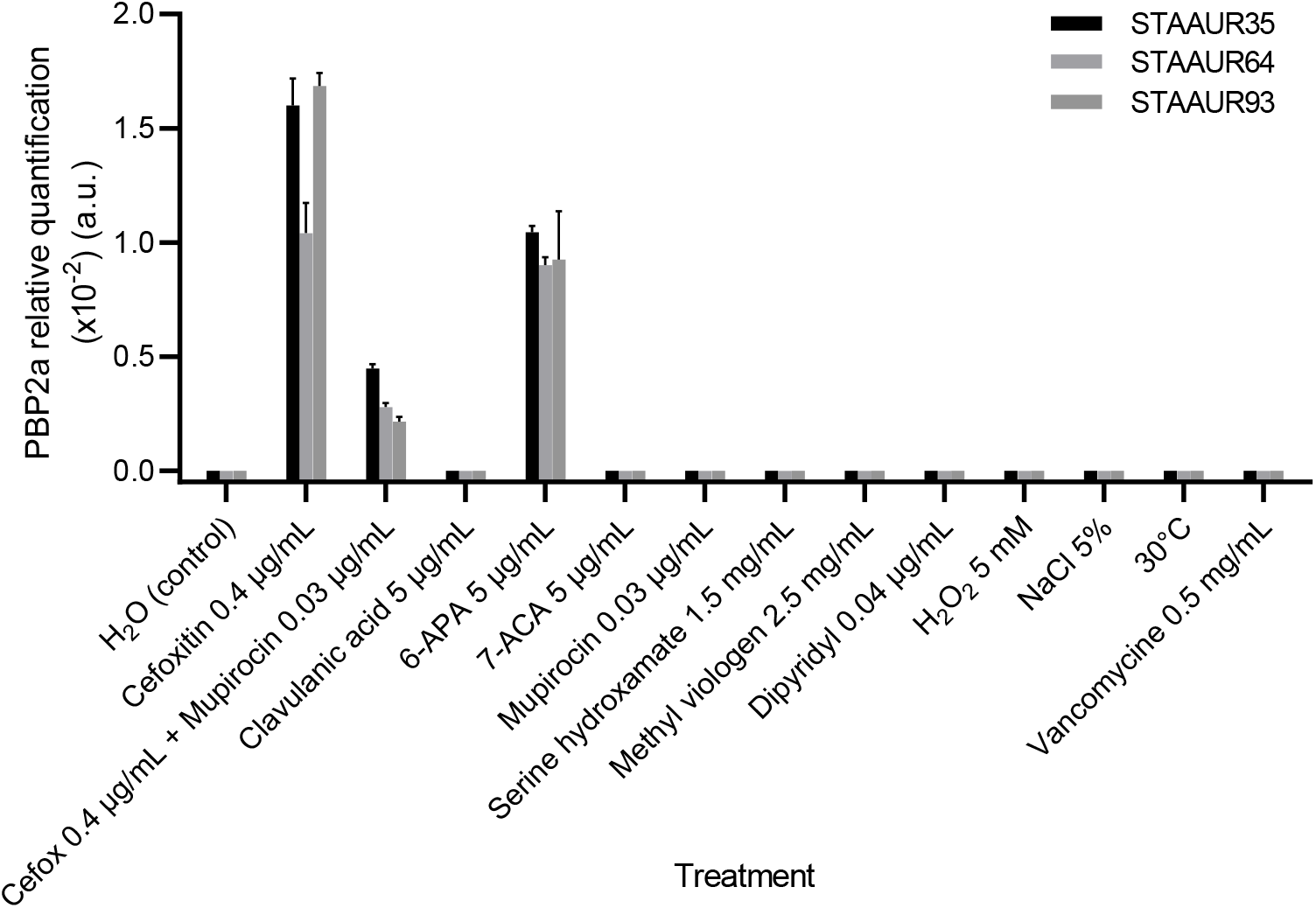
Evaluation of potential inducers to enhance PBP2a expression in 3 false negative MRSA strains after 1 hour of induction. Each condition was performed in biological triplicates (n=3).

Among all inducers tested, only cefoxitin, cefoxitin + mupirocin and 6-APA enabled PBP2a detection after a 1-hour induction. Surprisingly, PBP2a expression levels were lower when MRSA were treated using cefoxitin + mupirocin compared to cefoxitin alone (STAAUR35 p<0.05; STAAUR64 p<0.05; STAUR93 p<0.05; paired t test). Conversely, PBP2a peptides remained undetectable using clavulanic acid, 7-ACA, mupirocin, serine hydroxamate, H_2_O_2_, methyl viologen, NaCl, vancomycin, dipyridyl and incubation at 30°C. Compared to 6-ACA, higher detection levels were obtained using cefoxitin at a concentration of 0.4 µg/mL, which corresponds to a 1/10 cefoxitin clinical breakpoint (STAAUR35 p<0.05; STAAUR64 p=ns; STAUR93 p<0.05 (Figure 2).

To reduce the total turnaround time, duration of induction was further optimized for cefoxitin and 6-APA to enable a sufficient PBP2a detection with an acceptable increase of the total sample preparation time. The chromatograms and relative quantification presented in figure 3 show a clear time-response relationship of PBP2a expression upon cefoxitin and 6-APA induction, with a maximum being reached after 2 hours. The best compromise between time and quantifier peptide ELSISEDYIK relative signal was found to be 30 minutes of cefoxitin induction, as it was the shorter time required to detect PBP2a peptide in all three strains (Figure 3).

**Figure 3.**
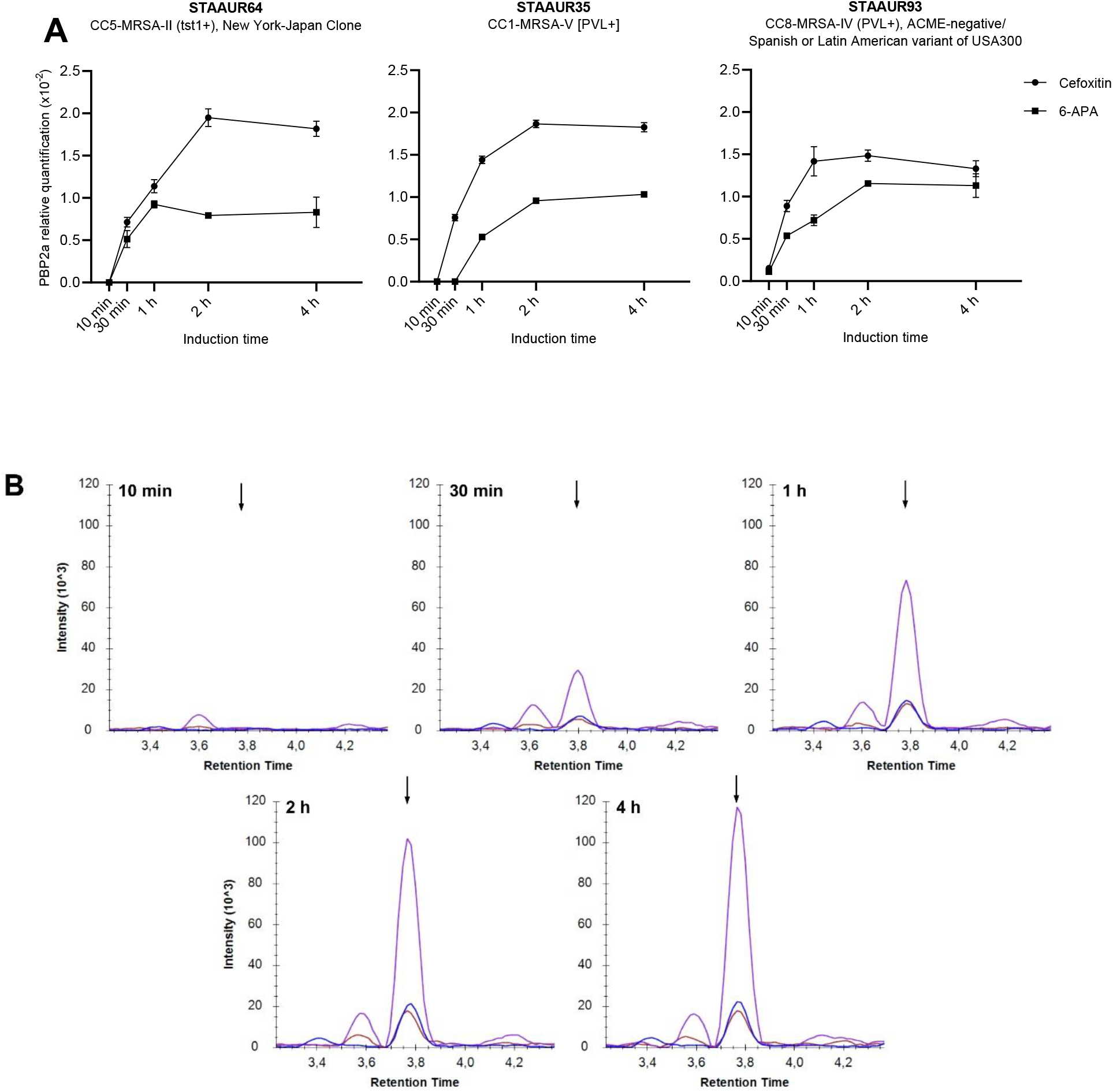
Kinetic of PBP2a expression upon cefoxitin induction. Strains were grown in blood culture bottles and induced with cefoxitin or 6-APA for 10 minutes, 30 minutes, 1 hour, 2 hours, and 4 hours after positivity. Each condition was performed in triplicate (n=3) **(A)** PBP2a relative quantification at different timepoints after cefoxitin or 6-APA induction **(B)** Reconstructed chromatograms for the ELSISEDYIK peptide (arrow) in strain STAAUR64 at different timepoints after cefoxitin induction. Purple, blue and red chromatograms correspond to the y6, y7 and y5 ions respectively.

### AME peptides selection

LC-MS/MS analysis of aminoglycoside-resistant strains carrying the corresponding AME genes as well as susceptible strains as negative controls led us to identify specific peptides usable for the detection of APH(3’)-III, ANT(4’)-I and AAC(6’)-APH(2’’). Peptide proteotypicity was confirmed by BLAST searching and signal specificity was verified with the corresponding synthetic peptides as illustrated in figure 4 for peptide AIGVYGSLGR of ANT(4’)-I.

**Figure 4.**
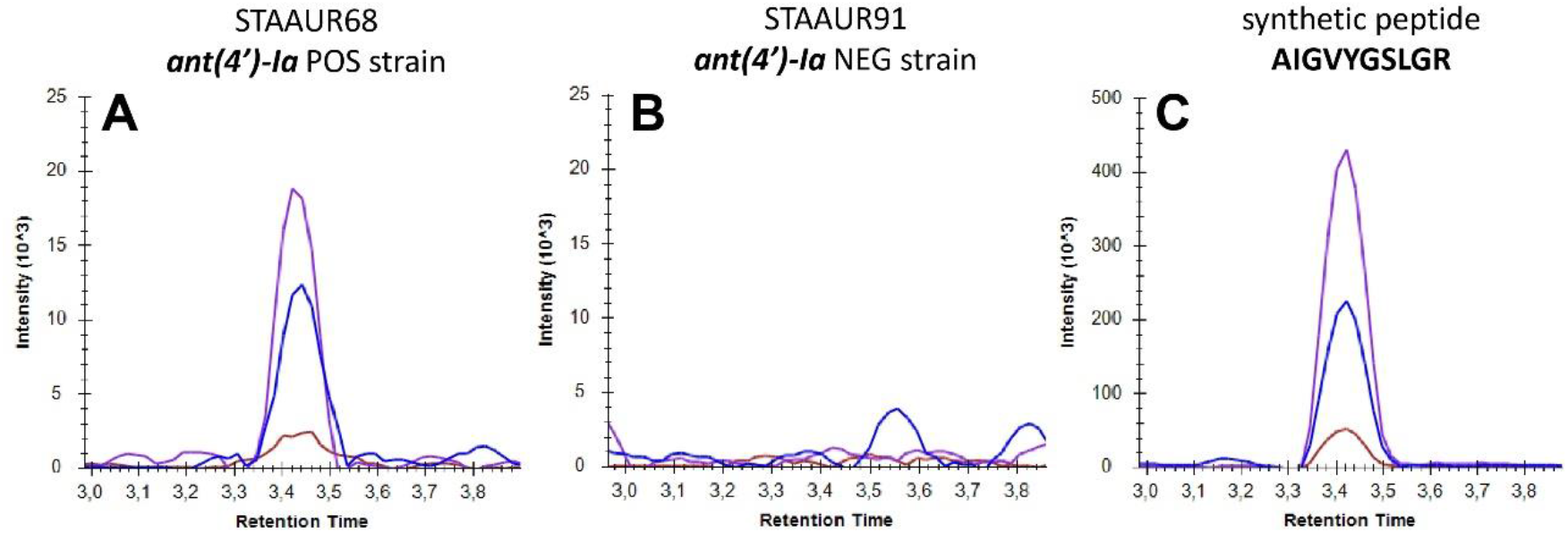
Specificity validation of ANT(4’)-I surrogate peptide AIGVYGSLGR. Signal observed in strain STAAUR68 (**A**) corresponds to the signal detected for the synthetic peptide (**C**). No interferences are observed in the ant(4’)-Ia negative strain STAAUR91 (**B**), confirming specificity of AIGVYGSLGR.

For each AME, the 3 best surrogate peptides were included in the assay to limit the total number of peptides monitored in the MRM method. AME surrogate peptides are listed in table 1.

**Table 1.**
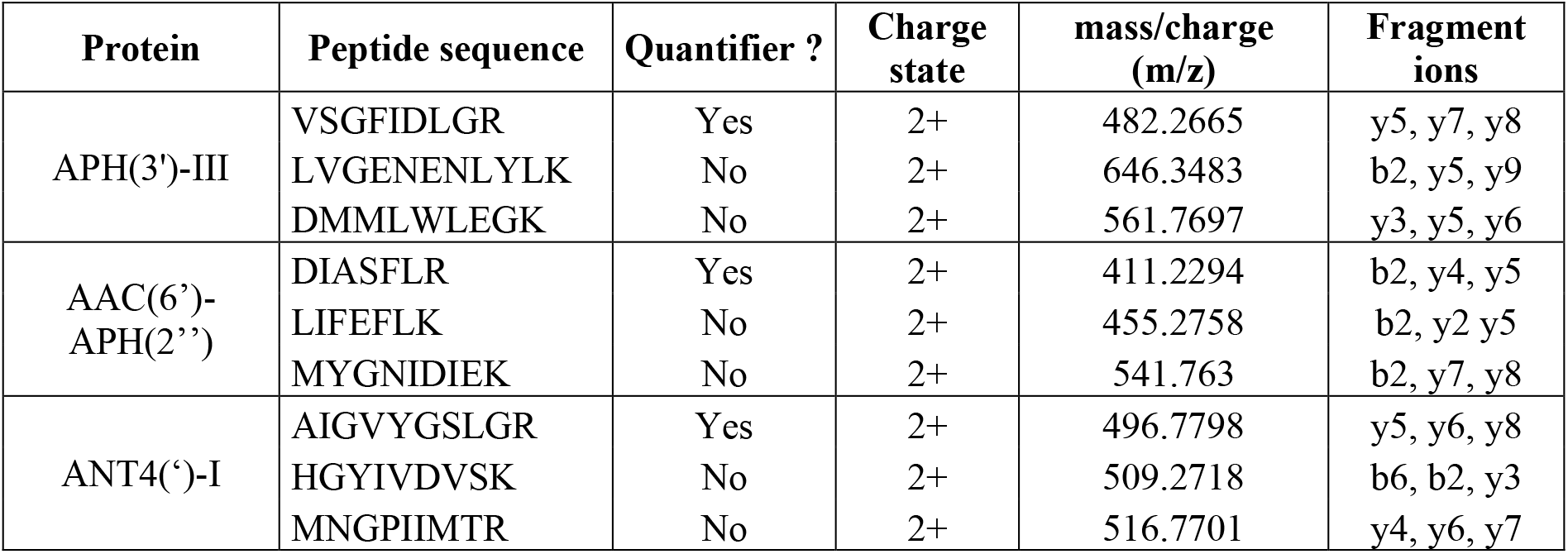
Surrogate peptides and fragment ions selected for detection of AME by targeted LC-MS/MS.

### Assay validation

A final 10-minute long LC-MS/MS method was setup targeting a total of 27 peptides; 17 resistance peptides (8 PBP peptides and 9 AME peptides), 6 ribosomal peptides, and 4 quality control peptides (Supplementary table S3). All 124 *S. aureus* strains were grown in aerobic and anaerobic blood culture bottles and underwent the 30-minute cefoxitin induction step upon positivity. The total time from blood culture positivity to the end of the LC-MS/MS run (cefoxitin induction, bacterial isolation from blood culture medium, bacterial lysis, protein digestion and analysis) was below 70 minutes.

To validate the analysis conformity, at least 3 *S. aureus* specific ribosomal peptides had to be detected as well as all quality control peptides. Overall, quality control peptides were detected in every sample along with at least 5 ribosomal peptides. The coefficient of variation (CV) of the retention time across the entire collection analysis was below 2% for every quality control peptide. Also, the CVs of one-to-one ratios of the three ribosomal quantifiers peptides were below 25%, confirming a satisfying repeatability of the trypsin digestion step (Supplementary figure S1)

Proteomic detection, WGS and AST results that were used to assess the sensitivity and specificity of the LC-MS/MS assay are detailed for each strain in the supplementary table S1.

Performances of resistance mechanisms detection are presented in table 2. Overall, the detection of resistance proteins was consistent between aerobic and anaerobic cultures, except for the STAAUR62 strain in which AAC(6’)-APH(2’’) was detected in the aerobic but not in the anaerobic sample, and for the STAAUR36 strain in which PBP2a was only detected in anaerobic conditions. Both inconsistencies can be explained by the very low expression levels of those proteins in these strains.

**Table 2.**
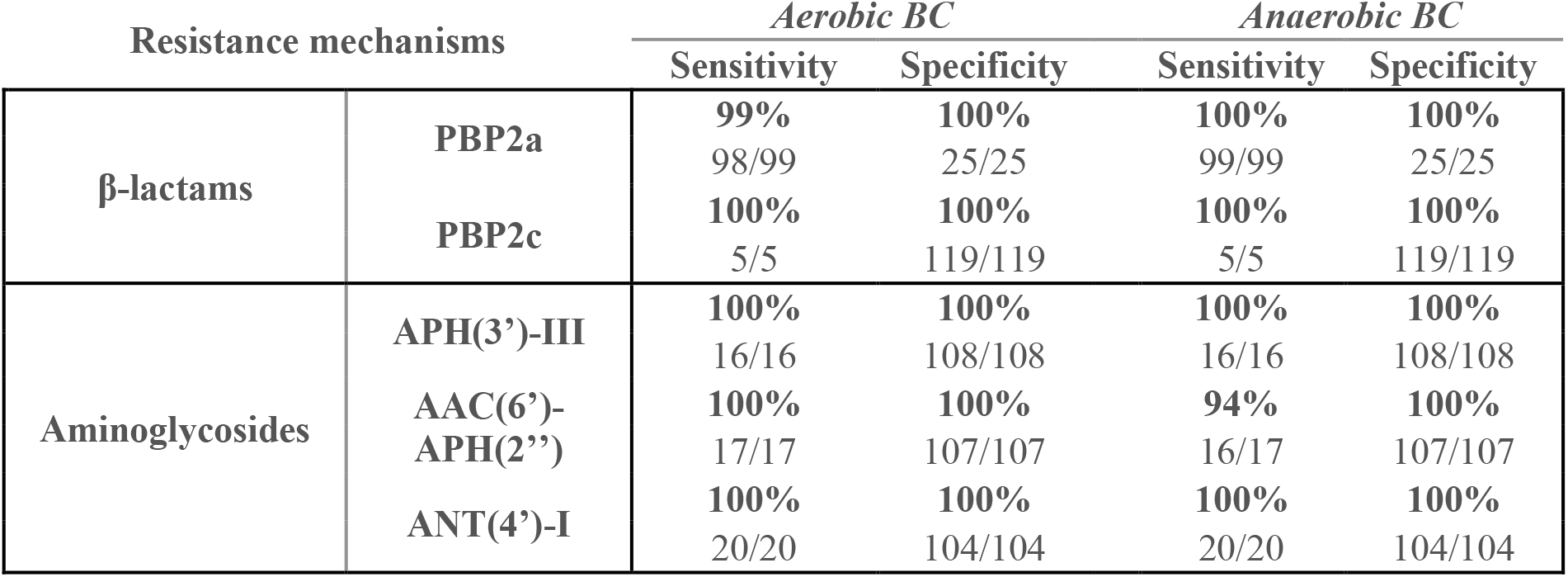
Performances of the multiplexed MRM assay targeting beta-lactam and aminoglycoside resistance proteins in 124 S. aureus strains directly from blood cultures compared to WGS data.

For PBP2c, APH(3’)-III and ANT(4’)-I, the sensitivity and specificity were both 100% compared to WGS. PBP2a was correctly detected in 98/99 aerobic samples and 99/99 anaerobic samples. Finally, AAC(6’)-APH(2’’) detection was consistent with WGS in 100% of the 17 aerobic samples but the detection failed in one anaerobic sample. For every resistance mechanism the specificity was 100%, given that no false positive results were observed compared to WGS.

Except for the STAAUR36 strain in aerobic condition, the present data show a total agreement between the presence of the *mecA/mecC* genes, the proteomic detection of PBP2a/PBP2c, and the phenotypic resistance to cefoxitin (Supplementary Table S1). This demonstrates that the LC-MS/MS approach can reliably characterize MRSA strains with performances comparable to the current gold standards (e.g. PCR for detection of *mecA* and *mecC*).

Similar results were observed for APH(3’)-III and the bifunctional enzyme AAC(6’)-APH(2’’), as respectively 16/16 and 16/17 strains showed consistency between genomic, proteomic and phenotypic resistance profiles for kanamycin, tobramycin and gentamicin (Supplementary Table S1). Relative quantification results are provided in the supplementary figures S2 and figure S3.

As for other resistance mechanisms, proteomic detection of ANT(4’)-I was in total agreement with the presence of the *ant(4’)-Ia* gene and the resistance to tobramycin (Supplementary Table S1). ANT(4’)-I is known to confer resistance to kanamycin but, using AST, the present data shows that among strains carrying *ant(4’)-Ia* as the exclusive aminoglycoside resistance gene, only 9 out of 19 strains were kanamycin-resistant (one strains was excluded because carrying the *aph(3’)-IIIa* gene in addition to *ant(4’)-Ia*). To further investigate this discrepancy, the relative quantification of ANT(4’)-I expression was performed for those strains (figure 5).

**Figure 5.**
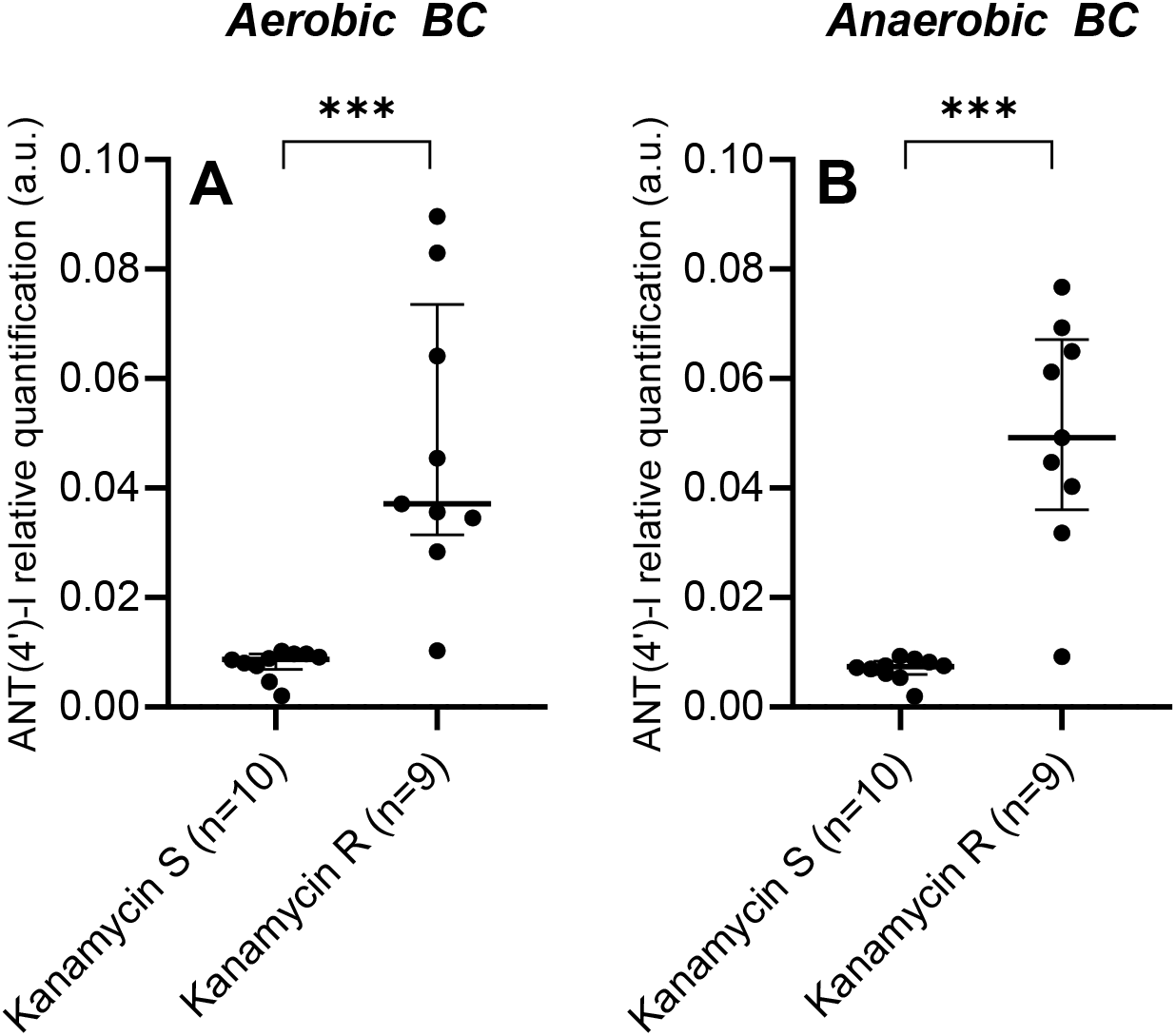
ANT(4’)-I relative expression in strains harboring ant(4’)-Ia as the only aminoglycoside resistance gene. Strains were cultured in aerobic (A) and anaerobic (B) blood cultures bottles. Strains are grouped according to their kanamycin resistance phenotype. S: susceptible; R: resistant. Statistical analysis was performed using a Mann-Whitney U test (***p<0.001). Horizontal bar represent the median and error bars represent the interquartile range.

The relative quantification revealed that ANT(4’)-I expression was significantly higher in kanamycin-resistant strains compared to the kanamycin-susceptible strains in both aerobic and anaerobic conditions. These data indicate that the presence of the *ant(4’)-Ia* gene alone is not sufficient to confer kanamycin resistance.

## Discussion

The present study reported a targeted proteomics workflow for the characterization of *S. aureus* resistance to beta-lactams and aminoglycosides in blood cultures, with a turnaround time below 70 minutes. A preliminary analysis of 99 *mecA* positive MRSA, representative of the worldwide SCC*mec* types diversity, allowed to unveil a broad heterogeneity in PBP2a expression levels between strains and an overall 43.4% sensitivity for PBP2a detection.

Several regulatory mechanisms are now well described, such as the MecI/MecR and BlaI/BlaR sensor/repressor systems that are directly involved in the modulation of beta-lactam resistance (43–45). However, it is unexpected to observe such differences in PBP2a expression between strains harboring similar SCC*mec* elements and MecI/MecR BlaI/BlaR regulons. These observations suggest the involvement of other regulatory elements such as two-component systems or global regulators (46, 47).

As a sensitivity close to 100% is expected for any diagnosis method, the induction was thus considered relevant to reach such levels of sensitivity. Cefoxitin is already used as an inducer to improve sensitivity of immunological test for PBP2a and PBP2c proteins detection (36). As the MecI/MecR and BlaI/BlaR systems are activated in response to beta-lactam exposure, clavulanic acid was also evaluated, as well as 6-aminopenicillanic acid and 7-aminocephalosporanic acid, two precursors in the synthesis of beta-lactam antibiotics (48, 49). It has also been shown that stringent stress, induced by amino acid deprivation after treatment with mupirocin or serine hydroxamate, led to increased PBP2a levels, particularly when combined with oxacillin (37, 38). Using the present culture and short induction conditions, no increase of PBP2a expression was measured. Surprisingly, mupirocin even had a negative effect on the inducing capacity of cefoxitin. Finally, H_2_O_2_, methyl viologen, NaCl, vancomycin, dipyridyl, and 30°C cold shock also had no effect on PBP2a levels, despite the evidences found in the literature (39–42). However, these evidences were based on prolonged exposure to the inducer, contrasting with the 1 hour exposure performed in the present study. This suggests a longer response time of the regulatory mechanisms activated by inducers other than cefoxitin and 6-APA. The 30-minute cefoxitin induction protocol allowed to achieve a 99% sensitivity and a 100% specificity for PBP2a detection, in both aerobic and anaerobic blood culture bottles. PBP2a was only missed in the aerobic sample of a ST59/952-MRSA-V(T) [PVL+] Taiwan Clone strain (among the 3 tested); a feature not typically reported in type-V SCC*mec* bearing strains (50, 51).

Gentamicin being a treatment of choice in *S. aureus* bloodstream infections, the detection of AAC(6’)-APH(2’’) was of prime necessity to provide a relevant diagnostic tool. Also, we observed that ANT(4’)-I detection *per se* only was not a good indicator to accurately characterize aminoglycoside resistance; a quantitative approach enabled to highlight a link between kanamycin MIC and ANT(4’)-I expression levels, and to distinguish between strains phenotypically susceptible and resistant to kanamycin. However, one strain among those with *ant(4’)-Ia* as the only aminoglycoside resistance gene showed low ANT(4’)-I expression levels, despite being phenotypically resistant to kanamycin. This suggests that an unidentified mechanism, such as ribosome mutation or modification/protection, may be responsible for this phenotype, as already described in Gram negative bacteria (52–55).

Moreover, it has to be mentioned that kanamycin is rarely used to treat *S. aureus* bloodstream infections, but the evaluation of kanamycin susceptibility serves as a proxy for amikacin treatment decision. Additionally, in clinic, ANT(4’)-I expression is deduced from a tobramycin-resistant profile, which also automatically classifies a strain as kanamycin-resistant. In the present study, all *ant(4’)-Ia* strains were phenotypically tobramycin-resistant, but not all were kanamycin-resistant. Our results suggest that the ANT(4’)-I-induced resistance is more complex than initially thought, which opens discussion on whether tobramycin susceptibility is the right indicator for ANT(4’)-I activity, or if amikacin could be used on tobramycin-resistant/kanamycin-susceptible strains.

One limiting factor to implement a new diagnostic tool in a clinical environment is the complexity of the sample preparation and the data analysis. Herein, to offer a rapid solution according to the need related to the diagnostic, the reduction/alkylation and solid-phase extraction steps, usually performed in proteomic analysis, were eliminated; and trypsin digestion, usually carried out overnight in conventional proteomic workflow, has been reduced to 10 minutes. Regarding data processing and output, an automatic system for chromatographic peak integration, signal validation, and clinical interpretation is currently being developed to provide easily readable information that can be interpreted by any hospital staff without chromatography and mass spectrometry expertise.

Among other factors influencing the introduction of new diagnostic tools in hospitals, the price is one of the most limiting parameters and directly depends on the cost of the reagents, the instrument, and the technical hands-on-time. Conversely to molecular tests such as PCR, the cost of reagents used in LC-MS/MS analysis falls below few euros per sample, and the major investment lies in the acquisition of the LC-MS/MS instrument. However, as documented by our group, the same LC-MS/MS platform can offer pathogen identification of species not differentiated by MALDI-TOF, identification of pathogen in polymicrobial samples, or detection of resistance mechanisms in other species (56, 57). Additionally, such assay can easily be updated according to epidemiological situations. The possibility to add new peptides to the analysis method, without the need to develop new reagents or to modify the sample preparation, enables an immediate response to the occurrence of new variants or resistance mechanisms. To our knowledge, there is currently no commercial multiplexed solution offering PBP2a/PBP2c qualitative detection and relative quantification of aminoglycoside modifying enzymes in less than 70 minutes and directly from a positive blood culture.

To conclude, this study presented an efficient targeted proteomic assay that enabled the prediction of *S. aureus* beta-lactams and aminoglycosides resistance directly from positive blood culture and with a turnaround time comparable to rapid molecular assays. Assuming an automated sample preparation and data processing, such method can have a significant impact on bloodstream infections diagnosis and can greatly improve patient management.

## Data availability

MRM raw data and transition lists are available via PASSEL with the accession number PASS05841 (https://db.systemsbiology.net/sbeams/cgi/PeptideAtlas/PASS_View).

## Acknowledgements

We would like to thank anyone involved in the IdBIORIV project, whether in the Institut des Sciences Analytiques, in the Hospices Civils de Lyon, in the Centre National de Référence des Staphylocoques or in the Centre International de Recherche en Infectiologie. We also thank Sciex company for providing the technical support and the instruments used in the present work.

This study and the IdBIORIV project were supported financially by the French country through the Agence Nationale de la Recherche and the « investissement d’avenir » program (ANR-18-RHUS-0013)

## Supplementary material

**Table S1.** Refer to the excel file

**Table S2.**
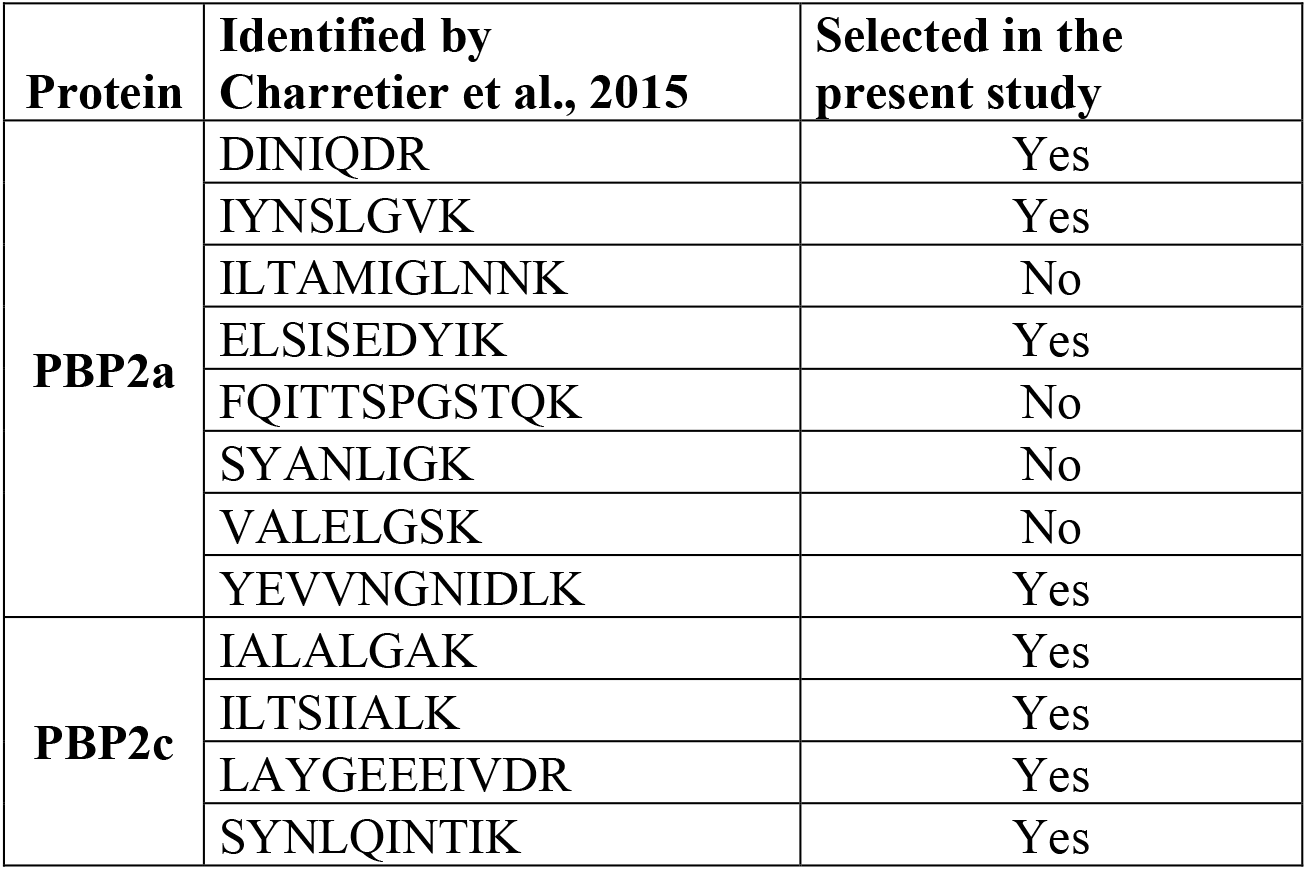
PBP2a and PBP2c surrogates peptides identified by Charretier *et al.*, 2015.

**Table S3.** Refer to the excel file

**Table S4.**
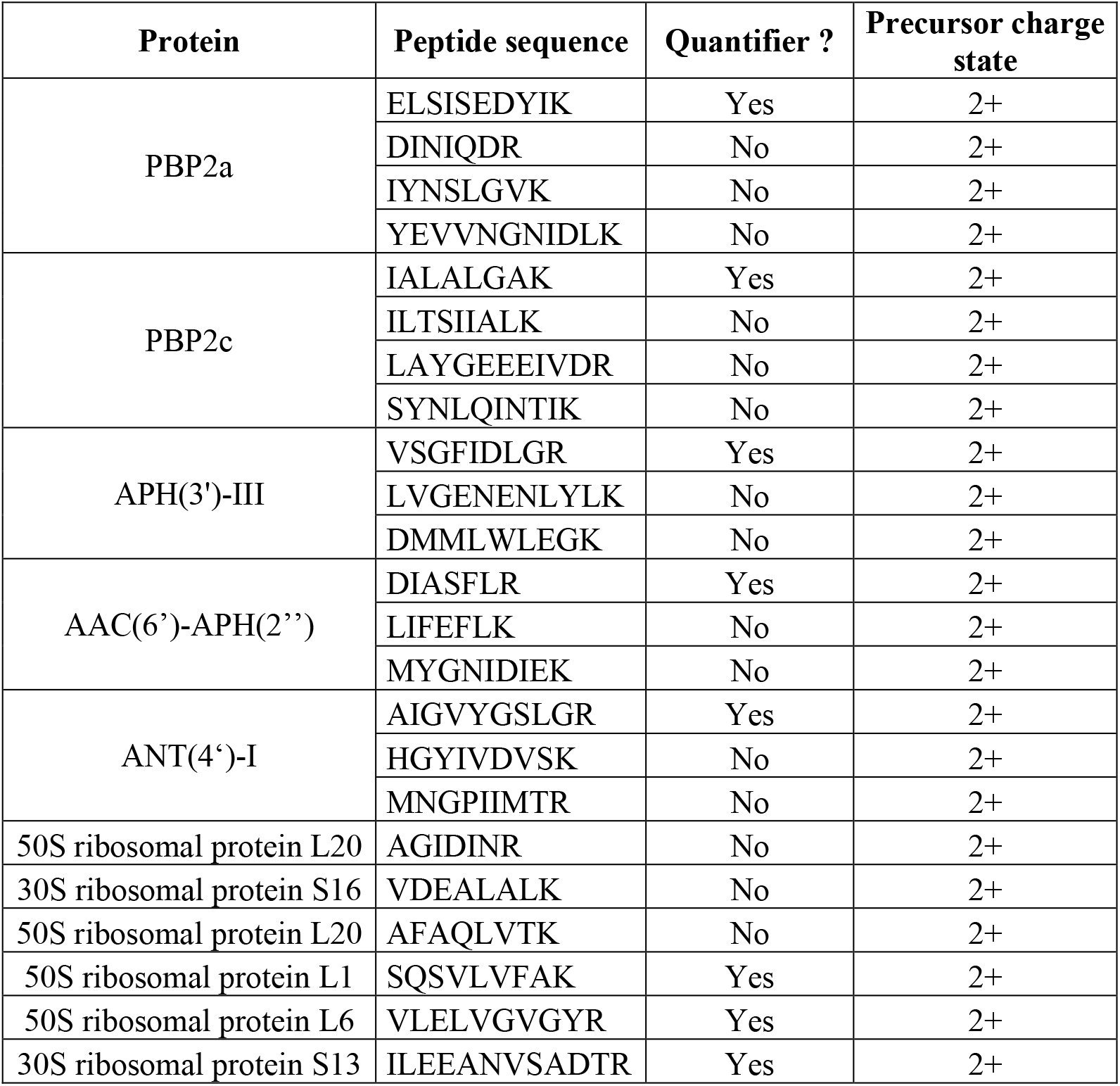
List of surrogate peptides used in the final targeted mass spectrometry assay.

| Protein | Peptide sequence | Quantifier ? | Precursor charge state |
| --- | --- | --- | --- |
| PBP2a | ELSISEDYIK | Yes | 2+ |
|  | DINIQDR | No | 2+ |
|  | IYNSLGVK | No | 2+ |
|  | YEVVNGNIDLK | No | 2+ |
| PBP2c | IALALGAK | Yes | 2+ |
|  | ILTSIIALK | No | 2+ |
|  | LAYGEEEEIVDR | No | 2+ |
|  | SYNLQINTIK | No | 2+ |
| APH(3')-III | VSGFIDLGR | Yes | 2+ |
|  | LVGENENLYLK | No | 2+ |
|  | DMMLWLEGK | No | 2+ |
| AAC(6')-APH(2'') | DIASFLR | Yes | 2+ |
|  | LIFEFLK | No | 2+ |
|  | MYGNIDIEK | No | 2+ |
| ANT(4')-I | AIGVYGSLGR | Yes | 2+ |
|  | HGYIVDVSK | No | 2+ |
|  | MNGPIIMTR | No | 2+ |
| 50S ribosomal protein L20 | AGIDINR | No | 2+ |
| 30S ribosomal protein S16 | VDEALALK | No | 2+ |
| 50S ribosomal protein L20 | AFAQLVTK | No | 2+ |
| 50S ribosomal protein L1 | SQSVLVFAK | Yes | 2+ |
| 50S ribosomal protein L6 | VLELVGVGYR | Yes | 2+ |
| 30S ribosomal protein S13 | ILEEANVSADTR | Yes | 2+ |

**Figure S1.**
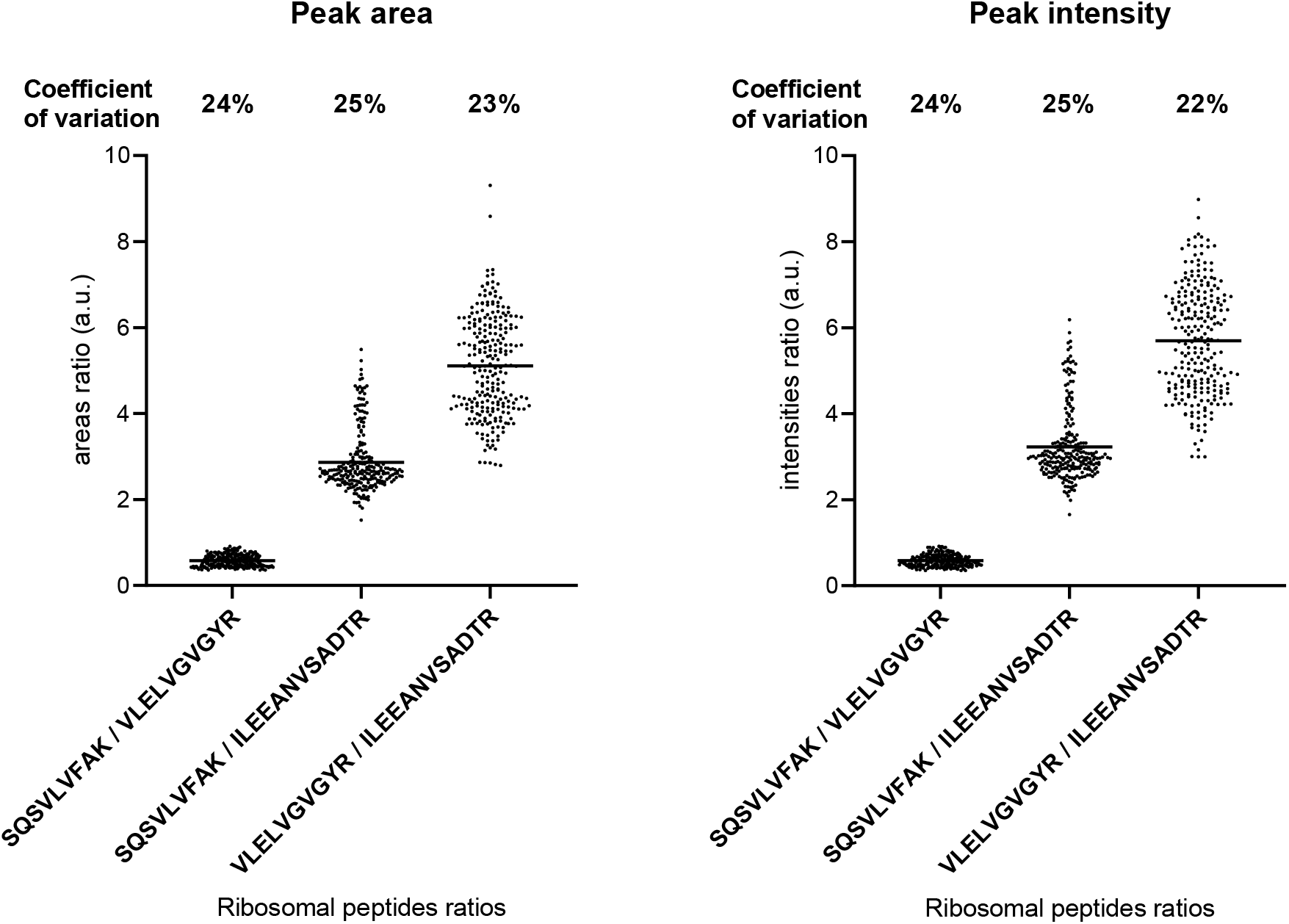
One-to-one ratios of the three quantifier ribosomal peptides. (A) Area ratios, (B) Intensity ratios. Those ratios are used to evaluate repeatability of the trypsin digestion step as relative intensities of ribosomal peptides should remain constant across samples.

**Figure S2.**
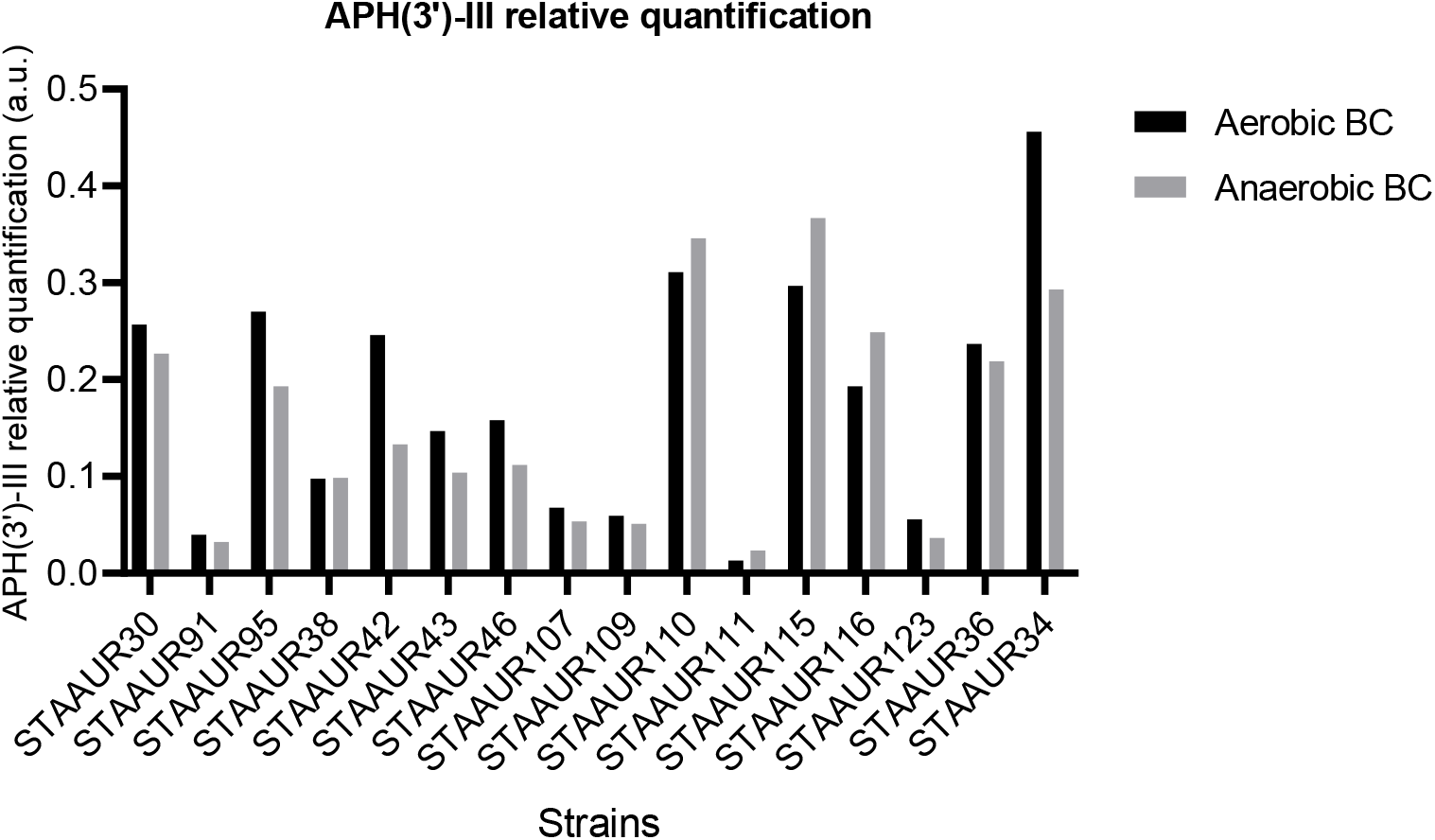
APH(3’)-III relative quantification in *aph(3’)-IIIa+* strains cultured in aerobic and anaerobic blood cultures. All strains are resistant to kanamycin.

**Figure S3.**
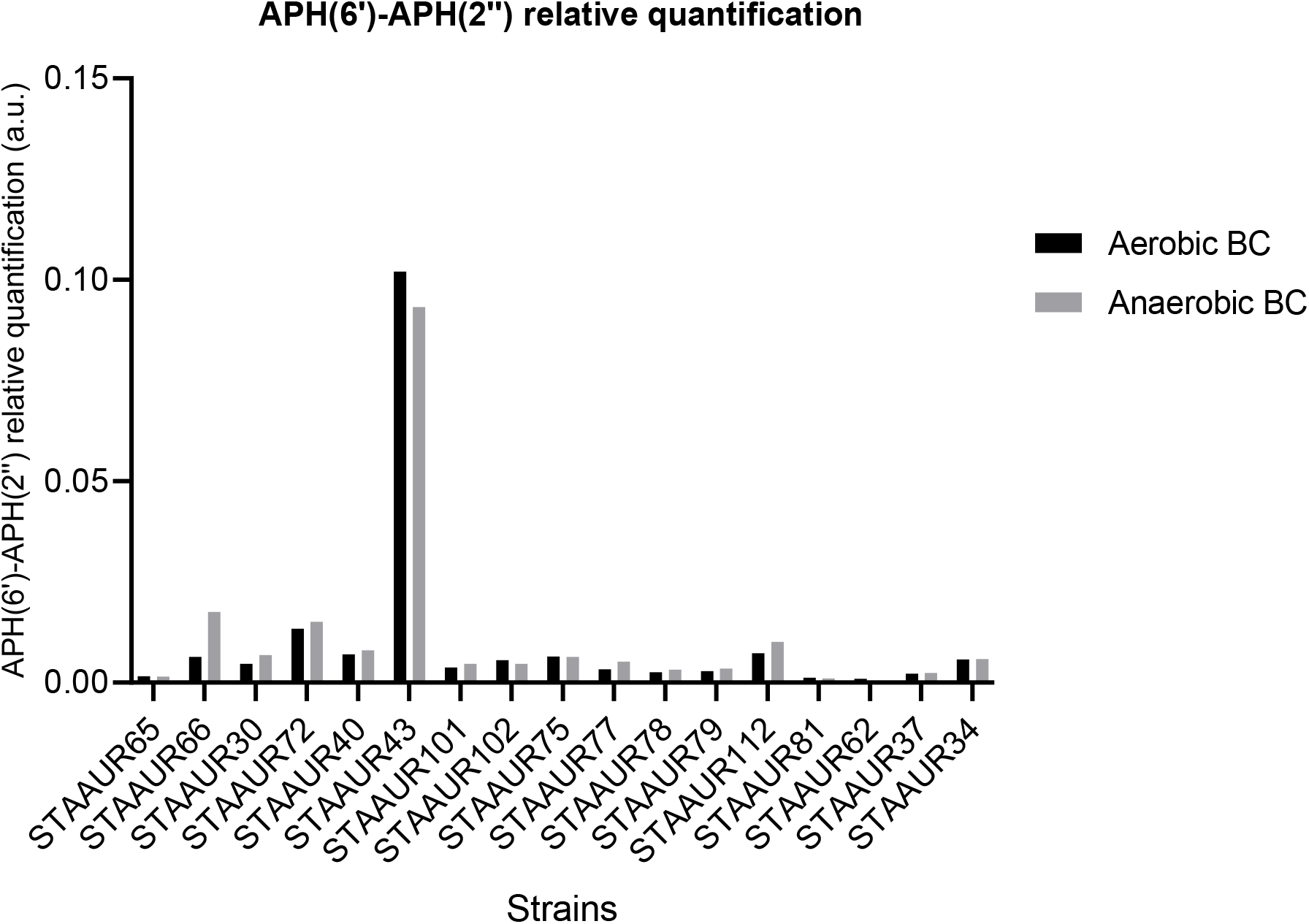
APH(6’)-APH(2’’) relative quantification in *aac(6’)-Ie-aph(2")-Ia*+ strains cultured in aerobic and anaerobic blood cultures. All strains are resistant to gentamicin. APH(6’)-APH(2’’)-I was not detected in the anaerobic sample for strains STAAUR62.

